# Collective flow of circadian clock information in honeybee colonies

**DOI:** 10.1101/2024.07.29.605620

**Authors:** Julia Mellert, Weronika Kłos, David M. Dormagen, Benjamin Wild, Adrian Zachariae, Michael L. Smith, C. Giovanni Galizia, Tim Landgraf

**Affiliations:** Department of Mathematics and Computer Science, Freie Universität Berlin, 14195 Berlin, Germany; Center for Digital Health, Berlin Institute of Health (BIH), Charite - University Medicine Berlin, Berlin, Germany; Center Synergy of Systems (SynoSys), Center for Interdisciplinary Digital Sciences, Technische Universität Dresden, Dresden, Germany; Department of Biology, Institute for Theoretical Biology, Humboldt-Universität zu Berlin, Berlin, Germany; Department of Biological Sciences, Auburn University, Auburn, AL 36849, USA; Department of Biology, University of Konstanz, 78464 Konstanz, Germany; Centre for the Advanced Study of Collective Behaviour, University of Konstanz, 78464 Konstanz, Germany

## Abstract

Honeybee colonies exhibit a collective circadian rhythm reflecting the periodic dynamics of the environment. Thousands of workers, including those engaged in in-hive tasks, must synchronize in various processes that may be rhythmic, such as nectar inflows, or non-rhythmic, such as brood care but it remains unknown how those different rhythms are integrated into a colony-level circadian rhythm. Using an AI-driven automated tracking system, we obtained uninterrupted long-term tracking of all individuals in two honeybee colonies. We demonstrate that circadian rhythmicity is present across all age groups and that this rhythm is entrained into all individuals, however, with peak activity shifting by up to 2 hours in workers furthest from the entrance. Extensive data analysis and an agent-based model suggest that mechanical interactions between individuals facilitate the transfer of movement speed, and hence Zeitgeber information. Finally, we show that this speed transfer leads to a collective slow wave of activity that initiates at the nest entrance, spreading throughout the nest. This simple mechanism, workers bumping into each other, enables colonies to entrain their rhythm to the daily cycle of the external environment and, because of the spatial organization of the nest, activates different groups of workers sequentially. The speed transfer interactions demonstrate a tightly-tuned mechanism that underlines the elegant self-organization of the superorganism.

## 2 Introduction

A molecular clock is present in nearly all living cells. For example, in the human body, all cells have a clock that is generated by a complex molecular feedback loop, involving, among others, the PER and CRY proteins [28]. Operating in rhythms allows cells to anticipate and respond to daily environmental changes, such as light and darkness. When each cell operates independently with its own unique rhythm and phase, it’s likely that the complex interplay of cellular processes would become disorganized. It is no surprise, then, that circadian phases in multicellular organisms are synchronized. Within the brain, for example, the suprachiasmatic nucleus of the hypothalamus acts as a major Zeitgeber for the remainder of the brain and the body [9]. Information about the daily rhythm from outside is provided by visual input from the eye, with both dedicated photoreceptors[1] and pooled information from all photoreceptors contributing information about light and dark day times. Information flow is provided by neurotransmitter release and hormonal regulations [9]. The coupling of multiple networks allows for the circadian clock in the mammalian body to remain synchronized across all tissues, with very little time lag. When this coupling is disrupted, leading to a desynchronization of peripheral and central clocks, the result leads to pathological situations and a diversity of diseases [9].

Social insects are classic examples of superorganisms: the reproductive unit is the entire colony, with hundreds to thousands of typically non-reproductive workers [12]. To operate as an integrated collective, however, the individual workers must coordinate their actions, similar to how cells unite to form a multicellular organism [27]. In colonies of the Western honey bee, Apis mellifera, different tasks are allocated via age polyethism and social experience: young individuals perform nursing duties, old individuals forage (to name but the two most prominent tasks; [20, 34]. Nursing duties are necessary throughout day and night, and are performed in constant darkness in the nest. In contrast, foraging duties are only possible with sufficient light during the day, and so foragers can rest at night [15, 14]. It is no surprise, then, that circadian rhythms have been shown in foragers, while nurses hardly show any circadian rhythm [7, 16, 29, 22, 23, 3]. Indeed, nurses have no access to external light sources and thus no direct abiotic circadian Zeitgeber. However, when removed from the nest, nurses do show a synchronized circadian rhythm, showing that circadian information is present, even when not displayed strongly in their behavior [10, 22, 23, 25]. These observations raise two important questions. First, how strict is the lack of circadian rhythm across nurse bees; do nurses and foragers form two independent groups, or is there a continuum that includes other castes/tasks? Second, is circadian entrainment independent across individuals in the colony and regulated by an abiotic Zeitgeber, or is it coordinated using a social Zeitgeber, and if so, what is the underlying mechanism? In the absence of light within the nest, abiotic common Zeitgebers could include temperature, vibrations or odor inflow. Conversely, a social Zeitgeber would rely on collective behaviors driving circadian synchrony. Previous data suggest that nurses are entrained by social factors such as volatile pheromones, vibration, and changes in the microenvironment such as humidity and CO2 concentration, though the details remain to be elucidated [10].

Here, we investigate the collective flow of circadian information within a honeybee colony, across all individuals. Recent developments in machine learning, tracking all individuals, and data analysis allowed us to follow the individual movement patterns of thousands of bees within the colony, over long periods of time and different years, for both foragers and nurses. We found that nurses, despite their round-the-clock work and lack of light exposure, do show a prominent (but weak) circadian rhythmicity. Furthermore, we find a shift in phase along a spatial gradient within the nest, suggesting that biotic rather than abiotic factors function as dominant Zeitgeber. Finally, we show that mechanical interaction among bees is a sufficient factor to explain the observed data: bees moving and bumping into each other act as social Zeitgebers to synchronize the entire colony into a common circadian rhythm. This is, to our knowledge, the first report about a collective system that entraines a circadian rhythm by mechanical interaction generating a sizable phase shift.

## 3 Results

We studied the age-related and spatial organization of circadian activity within honey bee colonies using long-term automated tracking of all individuals in two observation hives. Specifically, we continuously video-recorded two queenright colonies of Apis mellifera carnica over a period of 57 days (colony A) and 100 days (colony B). We regularly introduced uniquely marked bees, creating colonies in which each individual was identifiable, and of known age. The bees had access to the outside world via a tube, allowing them to forage in the wild, and were otherwise left undisturbed. Videos were processed to provide continuous trajectories and IDs for all individuals in the hive [32, 4, 35, 34]. In total, we tracked 1917 bees in colony A and 3404 bees in colony B. At any given day a minimum of 716 bees were visible (max: 1320 in colony A, 1594 colony B). We calculated each bee’s movement speed as the change of their position over time. Circadian rhythm was assessed for each bee by fitting a cosine with a period of 24 hours to movement speed data (Figure 1a). For each bee and each day, we used two parameters in subsequent analyses: the time of the activity peak *T* (*ϕ*) and the prominence of the rhythm (*R*^2^) (Figure 1b). For bee groups we calculated synchronicity S as the inverse normalized standard deviation of the phase (see Methods).

**Figure 1:**
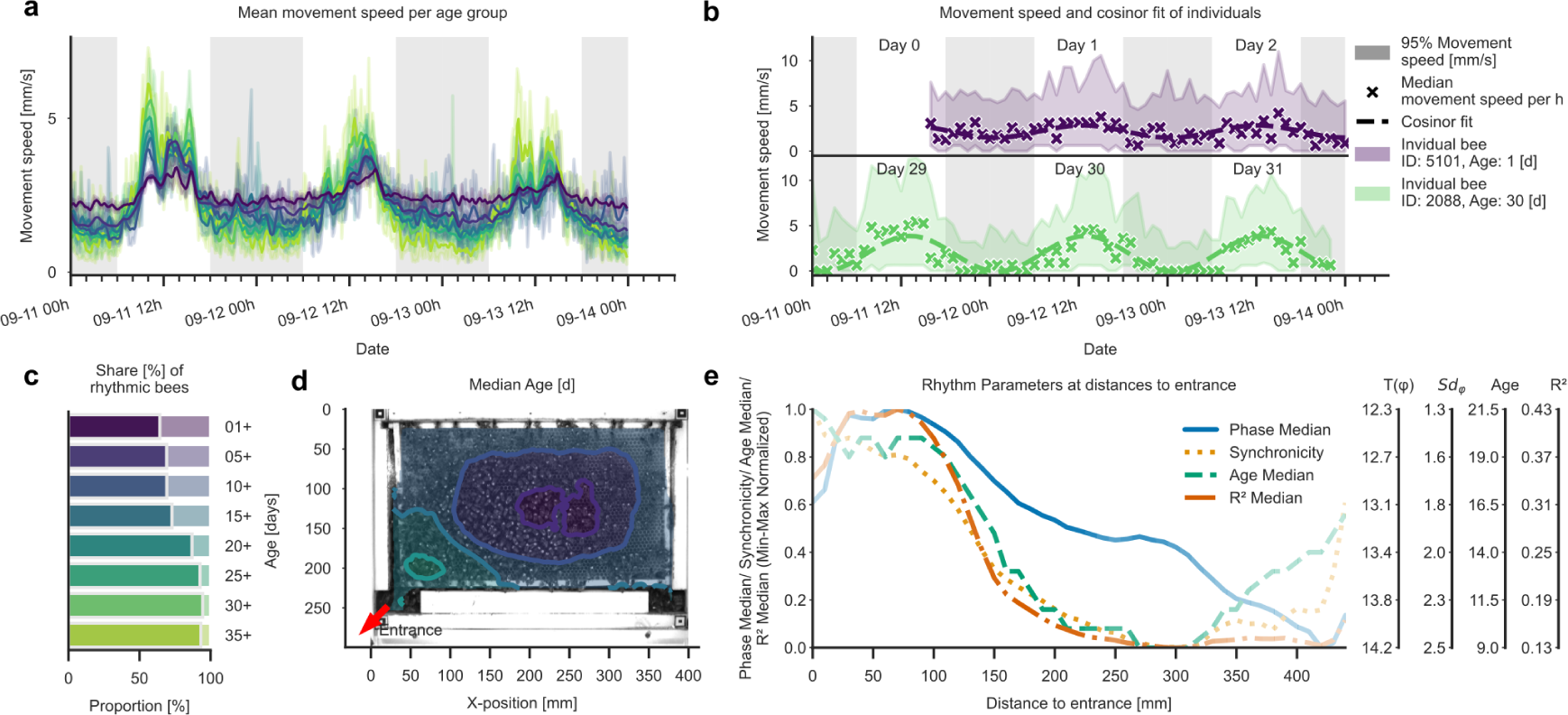
Circadian rhythms across bee age and nest space. **(a)** All age groups show higher speeds during the day than at night, but the difference increases with bee age. Example movement speed [mm/s] of bees for three days, split by age from young (purple) to old (yellow). Data was binned to hours, and smoothed with a Gaussian 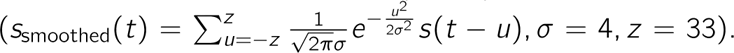 Lighter bands show 95% confidence intervals of the age group unsmoothed velocities. **(b)** Fitting a cosine function allows to extract circadian activity parameters. Two individuals are shown as example: a young nurse (top, purple) and an older forager (bottom, green), each for a three-day window. The cosinor fit is significant for both animals, but amplitude and variability are higher for the nurse bee. Note that some time points are missing for the young bee on day 0, since it was freshly introduced to the observation hive. **(c)** Spatial distribution of median age over the comb. Older bees (foragers) tend to be closer to the entrance, while younger bees (nurses) are more concentrated in the center of the comb. **(d)** With increasing age, the share of significantly circadian bees increases. Bees are split in age groups as in (a), and the proportion of significantly rhythmic bees is shown for each group. **(e)** Circadian parameters decrease with increasing distance from the entrance. Curves indicate peak phase *P* (daytime of maximum), phase standard deviation (hours), and circadian prominence *R*^2^ as calculated with the cosinor fit, relative to distance to entrance. Bee age (days) also decreases with increasing distance from the entrance. Curve color saturation indicates data density (i.e., how many bees were in that area). Synchronicity is defined as the standard deviation of the phase. The four scales to the right correspond to the original values for the four normalized traces. Data shown for colony B. For colony A refer to SI Fig. S1.

### 3.1 All ages rhythmic, distance to nest entrance linked to rhythmicity, synchronicity, and phase

First, we analyzed how circadian rhythmicity changes with an individual’s age. We split bees by age cohorts of 5 days each, and quantified the percentage of rhythmic bees within each group(Figure 1c). We found significant daily rhythms in all age groups, and with increasing age the proportion of rhythmic bees increased. For the youngest cohort (bees aged 1-4 days), we found 29% and 64% significantly rhythmic bees in colony A and B, respectively (for the very first day, the numbers were 18% and 58%, respectively). Conversely, more than 89% and 93% of the bees aged 35 days or more had significant circadian rhythms (Figure 1c).

Next, we investigated how age and rhythmic activity are organized spatially in the nest. To do this, we mapped the median age for each position on the comb. As expected [31, 21], we found a systematic and continuous decrease in age with increasing distance from the nest entrance (Figure 1d). This decrease in age is correlated with a delay in peak activity time, shifting by almost two hours, from 12:20 to 14:12 (right axis Figure 1e). Therefore, the further away a bee is from the nest entrance, the later her activity peaks in the day. Interestingly, rhythm prominence also decreases with increasing distance from the entrance (*R*^2^ in Figure 1e), and bees are also less synchronized (synchronicity in Figure 1e). This spatial pattern in rhythmicity is likely due to age polyethism: young bees are nurses caring for brood in the center of the nest, middle-aged bees offload foragers at the entrance and process nectar into honey, and foragers collect resources outside and pass them off to nestmates close to the nest entrance. Therefore, the spatial organization of circadian phase and activity strength within the nest can be mapped onto bee castes: nurse bees show a weaker rhythm and a later activity peak than foragers. The continuous character of this shift (Figure 1e) reflects the continuity of cast development across several tasks within the lifespan of bees [34]. Note that the distance relationship is not strictly monotonic: in Figure 1e, the youngest workers are 300 mm from the entrance (brood area), and with even further distance age increases again. Importantly, however, peak activity time further decreases with increasing distance to the entrance even beyond the brood area.

### 3.2 Local interactions yield speed transfers

How do bees inside the dark nest entrain onto a circadian rhythm? If the Zeitgeber was an abiotic factor, such as light diffusing from the entrance (instantaneous), or floral odors spreading (within milliseconds), or temperature [13], we might also expect a reduced prominence in circadian rhythm with increasing distance from the entrance, but we would expect the rhythm to be synchronized. Therefore, we hypothesized that circadian entrainment could be caused by physical interaction among bees. This would create a local mechanism that spreads activity from one individual to the next [30]. To test this idea, we identified all interactions of bees by quantifying their proximity: we defined each bee as a rectangle of 14 mm by 6 mm, and extracted all pairs with overlapping rectangles. For each interaction, we analyzed the speed change by subtracting the average speed of 30 seconds before an interaction from the speed 30 seconds after the interaction. Since this was done for each bee in a pair (every bee was a “focal bee” and an “interacting partner” once), each interaction yielded two speed-change data-points. We grouped both the focal bees and the interacting partners into six quantiles, and plotted the speed change for all pairings in a 6×6 matrix of 36 cells (Figure 2a, left panel). We found that fast bees become slower, in particular when interacting with slow bees, and slow bees increase their velocity upon interaction, in particular when interacting with fast bees (Figure 2a, left panel “Real data”). This indicates that physical interaction leads to a transfer of movement activity.

**Figure 2:**
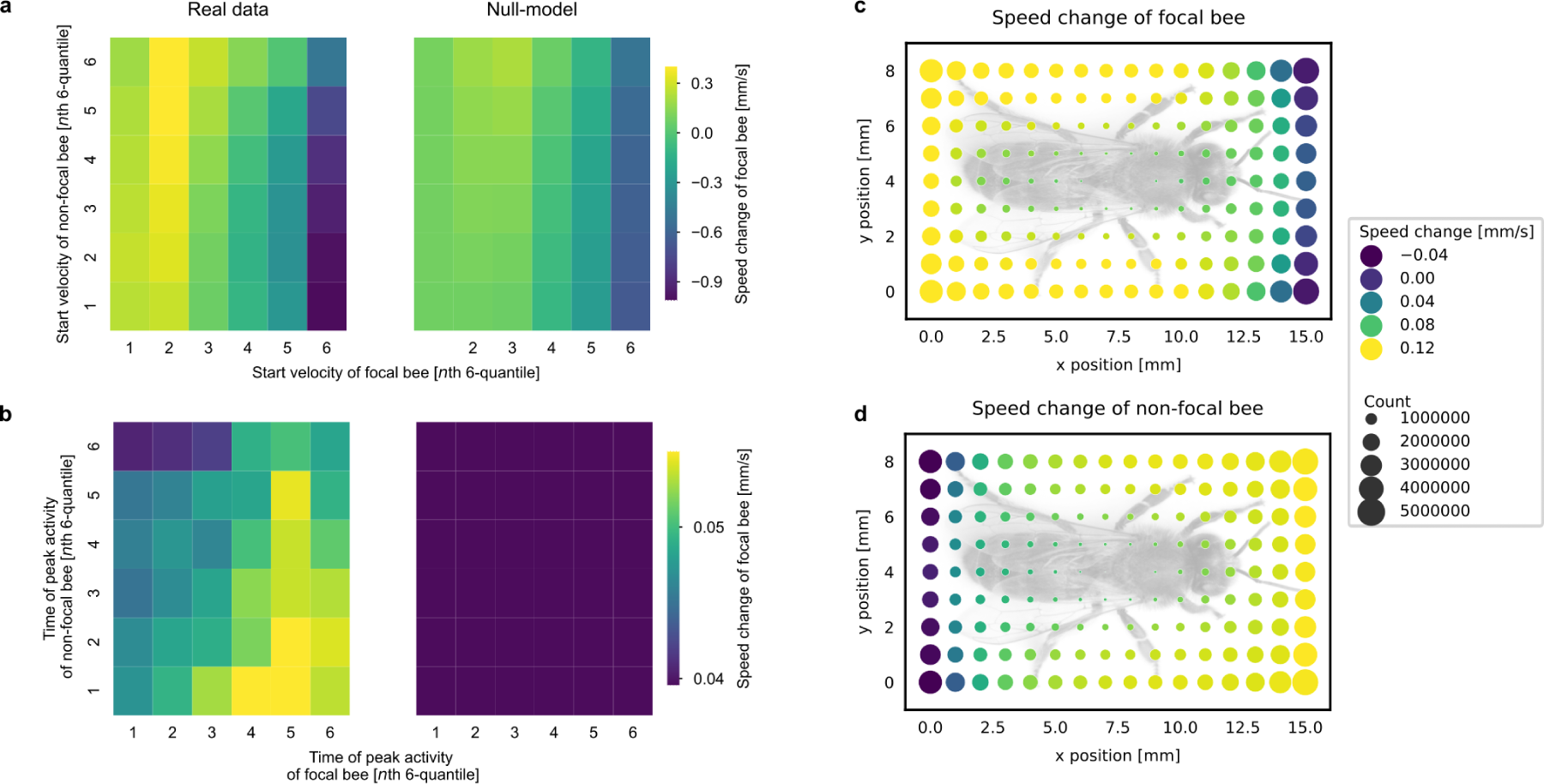
Dynamic physical interaction leads to speed transfer across bees. **(a)** Fast bees are slowed down when bumping into slow bees. Focal bees (x-axis) and non-focal bees (y-axis) were sorted into 6 speed quantiles. For each combination, the speed change of the focal bee was color coded: speed decreases (blue hues) were particularly prominent for previously fast bees (6th quantile). The right panel shows data from a randomly shuffled null-model (see methods), experimental data is significantly different from the shuffled distribution (see Supplementary Section 10.2 for p-values). **(b)** Circadian phase *ϕ* of the interaction partner influences speed transfer. Focal and non-focal bees were binned in 6 quantiles depending on the phase of their cosinor fitted circadian activities. Focal bees with late phase led to higher increases in non-focal bees with early phase, and focal bees with early phase led to small increases in non-focal bees with late phase. The randomly shuffled null-model (right panel) showed no phase effect. **(c)** Bees are slowed down when bumping into another bee from the front, and accelerated when bumped from the back. Image shows speed change of the focal bee (shown schematically in grey) relative to the interaction location along her body. Speed changes resulting from interacting at the front are negative (blue hue), interactions at the back are positive (yellow hue). **(d)** Bees are slowed down when bumping into the back of another bee, and accelerated when hit by a head. Image shows speed change of the non-focal bee relative to the interaction location along the body of the focal bee (shown schematically in grey). Speed changes are negative along the back of the focal bee, and positive along her front. In (c) and (d), circle sizes indicate the number of observations in each point.

Is this movement transfer significantly different from the general speed changes in the colony that are not linked to physical interactions? To test this, we constructed a null model by sampling two bees on the comb surface randomly for every interaction time point and assessing their speed changes as before. Even in this baseline condition, fast animals tend to become slower and slow bees on average become faster (Figure 2a, right panel, “Null model”). This can be explained, in that speed values are bounded. When a bee is moving at its maximum speed, any speed change can only be a deceleration due to energy constraints and mechanical limitations. By the same logic, bees that do not move can only remain still or accelerate. However, when we statistically tested the null-model against the real data, we found a significant difference in the vast majority of cells (Welch’s t-test p ¡ 0.05 in 35 out of 36 cells for colony B, 34 out of 36 cells in colony A). Social interactions specifically, thus, are a major cause for movement speed change. Analogous to physical particles transferring momentum when they collide, we can think of this process as a “speed transfer” between two individuals.

Grouping bees by time of peak activity *T* (*ϕ*), we find that bees with a late peak of activity get accelerated the most, in particular by early peaking bees which on average are older, closer to the entrance and more pronounced in their rhythm (Figure 2b, left panel). A similar observation is valid for age and *R*^2^ (see Supplementary Information S3). Again, a comparison with shuffled data in a null model (Figure 2b, right panel) shows a high significance (36 out of 36 with p ¡ 0.01 ??). Speed change for these groups is less pronounced as when comparing speed groups directly, suggesting that the driving force is speed itself, and not age, *R*^2^, or *T* (*ϕ*). The latter show a significant effect due to the inherent correlation of speed with age and or phase *ϕ*.

Speed transfer across bees might be due to a non-directional arousal effect, where being touched by another bee leads to increased activity, or it could be due to a directional nudging effect. To differentiate between these alternatives, we mapped speed changes as a function of where on the body the interaction took place, by projecting speed change onto the ego-centric coordinate system of each bee (here referred to as focal bee). Then, we plotted how the focal bee changes speed depending on where she touched her sister (Figure 2c). The resulting map shows that, on average, when a bee touches her sister with her head she is slowed down (blue circles in Figure 2c), while when she interacts at the sides, and more so at the back, she speeds up (yellow circles in Figure 2c). Conversely, her interaction partner becomes faster when being touched by the focal bee’s head (yellow circles in Figure 2d), and slows down when being touched by the focal bee’s back (blue circles in Figure 2d). Given that bees generally move forward, this result suggests a simple physical interpretation: when a bee bumps into the back of a nestmate with her head, she slows down, while her partner is accelerated. Speed transfer is a directional nudging effect. However, the situation is different for resting bees (speed ¡ 1 mm / sec): these bees accelerate with every interaction, suggesting that for resting bees an arousal effect, independent from the interaction site, is the best explanation (see Supplementary Figure S3).

### 3.3 Speed transfers cause a wave-like propagation of activity

The observed directional nudging effect leads to the hypothesis that young nurse bees are entrained into their circadian rhythm by the activity of older forager bees. Is this mechanism sufficient to explain the observed movement patterns in the nest? To test this hypothesis, we implemented an agent-based simulation in which two groups of agents perform random walks and transfer their movement energy when spatially interacting. The two groups start off with a spatial preference for a subregion of the nest. The members of one group move at a constant speed starting at the nest center (representing the young bees deep inside the nest), the others exhibit a sinusoidal speed fluctuation and start close to the entrance of the nest (representing the old bees that are synchronized to external conditions). Social interactions transfer speed similar to the effect observed in the data, such that slow bees gain speed from interactions. Similar to atoms at higher temperatures, the distribution of the simulated agents expands further into 2-D space when increasing their speed. The virtual nest is confined: agents moving into the walls are reflected back. With increasing simulation time, the rhythmic group fills the available space and the number of interactions with the non-rhythmic group increases. Due to these interactions, the non-rhythmic agents move faster, spread out further, and, in effect, interact more with other agents. These interactions, however, take time to build up, creating a pronounced phase-shift in the activity of both groups. This phase shift depends on factors that determine how much time passes between social interactions, such as the speed of movement, how much both groups overlap initially, how far apart the two preferred locations are, and the standard deviation of the gaussian used to draw the initial locations of rhythmic and non-rhythmic agents (data not shown). The agent-based model shows a phase shift with increasing distance to the rhythmic group, replicating the situation found in the experimental bee data, where the phase *ϕ* shifts with increasing distance to the nest entrance (Figure 3a,b). In fact, circadian activity, as a result of bumping physical interactions, creates a wave of increased activity moving from the entrance into the inner area of the comb. The simulation shows that the speed transfer model is sufficient to explain the spreading of rhythmicity and the shift of the activity phase. It should be noted, though, that the model does not exclude the existence of additional mechanisms in a real bee colony.

**Figure 3:**
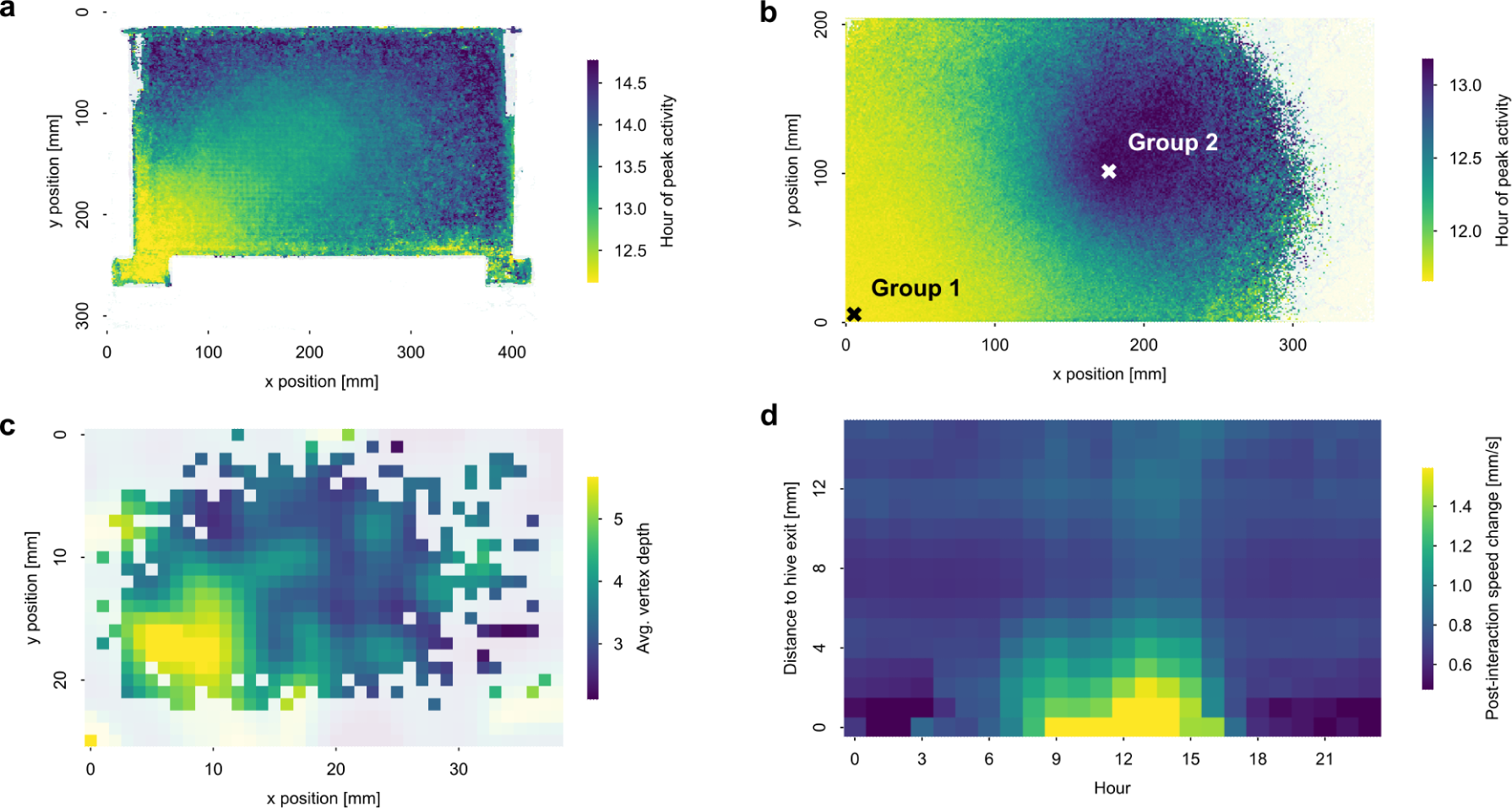
Activity propagates in a wave-like manner through the nest. **(a)** Peak activity time (phase *ϕ*) is early at the nest’s entrance, and increasingly late with distance to the entrance. Median phase values were color-coded onto a view on the comb. Entrance is at the lower left. **(b)** The agent-based model replicates the same spatial pattern of spatial phase distribution. Peak activity time (phase *ϕ*) was color-coded with a view on the simulated comb for the agent-based model. Crosses show centers of mass at the beginning of the simulation run for group 1 (initially rhythmic, lower left) and group 2 (initially non-rhythmic). **(c)** Bees initiating interaction trees are mostly localized at the nest entrance. The plot shows the average depth of interaction tree vertices at each nest location. The larger the depth, the more interactions will be downstream of that bee. **(d)** Largest interaction effects are localized at the entrance, and occur around noon. This two-dimensional false-color coded plot shows speed increase resulting from an interaction as a function of time-of-day (x-axis) and distance to the entrance (y-axis). The deeper into the nest, the weaker and later we found speed transfers.

If, as suggested by the simulation results, speed transfers are causing the non-rhythmic nurses to show rhythmicity, then it should be possible to track the initiating agents, when all bees are monitored over time, as is the case in our experiments. Therefore, we first identified the individuals that had an activating effect on a given young and rhythmic bee and recursively applied this procedure backwards in time. Every bee potentially had multiple activating interactions with multiple activating bees. Tracing back all activating paths, thus, yields a tree structure with the root representing a target nurse bee and a variable number of bumping bees per level that had speed-increasing interactions. The leaves of the resulting trees are the origins (driver bees) of activating chains of speed transfers ultimately entraining the focal bee at the root. We find that these driver bees exhibit a significantly earlier phase *ϕ*, higher rhythmicity *R*^2^, and are older than the remaining bees across the entire nest (Brunner-Munzel test, p ¡ 0.001). This is reflected in the observation that deeper trees (with more layers) are initiated closer to the nest entrance (Figure 3c). Thus, we have shown that speed transfers travel from older bees with highly pronounced rhythms close to the nest entrance to younger, less rhythmic bees further inside the nest.

We analyzed the spatio-temporal distribution of bumping speed transfers. We plotted speed increase (color coded) against time of day (x-axis) and distance to the entrance (y-axis), and show that the bumping speed transfer is strongest around mid-day (9 am to 3 pm), and close to the entrance (distance ¡ 15 cm) (Figure 3d). Activity and speed transfers propagate like a wave, from the entrance deeper into the nest. As the wave travels inwards, the speed changes diminish gradually. This corroborates our interpretation that activity levels propagate from the entrance to the deeper areas of the nest, and that the entrance to the nest represents the origin of bumping interactions and of activating cascades eventually giving rise to the rhythmicity of the young bees in the brood nest.

## 4 Discussion

Prior work on circadian rhythms in foragers visiting flowers [31], or arrhythmic nurses performing round-the-clock brood care [16, 3], have examined individuals, but colonies are integrated collectives. Therefore, we examined how workers coordinate their temporal activity rhythms within the collective. Taking advantage of recently developed tracking technology [26, 35, 4, 34] we mapped the movement of all bees within the hive, without interruptions, across multiple days, and with known identity and age for each individual. We found that all bee casts display circadian activity, but nurses do so with less intensity. We also found that circadian activity is synchronized across bees through mechanical interaction, leading to a 2-hour time shift in activity peaks within the hive.

Interestingly, we did not find a clear distinction between “nurses” as a group, and “foragers” as another group. Rather, we found a continuum: the oldest bees, most of them foragers who often left the hive and were found close to the hive’s entrance, had a strong circadian rhythm. The distance from the nest entrance predicts key properties of the daily rhythm: the further from the entrance, the weaker the rhythm, the lower the synchronization between bees, and the later the activity peaks (Figure 1). Thus, bees without access to external circadian cues are still synchronized to day and night cycles. This suggests that circadian behavior is not dictated by a particular task, since in that case we would have expected distinct groups of bees displaying a particular phenotype. Rather, circadian behavior is best explained by spatial location within the hive.

We identified cascades of mechanical speed-transferring interactions as the main mechanism driving this rhythm. Circadian timing emerges as a property coordinated via local arousals propagated throughout the nest. This behavioral mechanism exploits the individual circadian propensity of each bee: the bumping does not create a circadian rhythm, but rather only entrains the phase of the rhythm. As a result, we observed a wave of increasing activity traveling across the comb.

This wavelike phenomenon is not a transfer of mechanical energy but can be thought of as an information wave that changes the statistical properties of the individuals it reaches. Unlike in multicellular organisms where hormones and/or neural signals create almost instant synchrony [9], the speed transfer cascades produce activity in deeper parts of the nest with a time-lag of up to two hours. Given the time required to find and bring in food, this lag appears to be adaptive: food-processing bees become active only after most foragers are already busy. The speed transfers have an additional benefit. Where individuals move a lot, passage for others becomes more permissive. Thus, when the nest becomes busier and many workers engage in various tasks, moving resources and information becomes more effective. This positive feedback loop (more active workers, more speed transfers, more movement energy) allows for activating nest regions and optimizing movement throughput.

Our data cannot exclude that, in addition to the behavioral ‘bumping’ mechanism, other Zeitgebers might contribute to synchronizing bee groups. It has been shown that social factors such as odors from the outside world brought in by foraging bees, vibrations (both from outside or caused by returning foragers dancing on the comb) [17, 24, 25]. Abiotic factors such as temperature shifts and diffusing light might also contribute, but are not necessary [25]. However, all of these factors have in common that their spatial spread is fast, if not immediate. Indeed, their effect was studied using individuals removed from the hive. Thanks to our approach to observe all bees, over long time periods, within a full hive, we were able to demonstrate a 2-hours time lag, which is a strong indication that the dominant mechanism for social entrainment is a mechanism that is slow and behavioral. Thus, we propose that mechanical bee-to-bee interaction is the main synchronizing mechanism across bees.

The superorganism honeybee colony has often been compared to a multicellular organism. Here we show that, for the organization of circadian timing, the bee superorganism uses a slow spatial mechanism unknown in multicellular organisms. While in, say, the mammalian body circadian rhythm is synchronized using hormones and neural signals [9, 19], the behavioral bumping mechanism in bees allows for an additional level of complexity: the activity peak shifts within the hive, in accordance to the shifted need for an activity peak for each task. While foragers have to collect nectar and pollen first, receiving bees have to become active only after foragers come back from their first flight in the day, Accordingly, honey processing bees enter their chores only later. We are not aware of any cell-to-cell mechanism across neighboring cells in multicellular animals being used for synchronizing circadian activity and creating a relevant time-shift. Thus, the superorganism “honeybee colony” has evolved a new strategy for circadian synchronization, unknown to multicellular organisms.

Taken together, we show that a bee hive as a collective is photically entrained by external Zeitgebers sensed by the forager bees, who have access to the day/night light cycle. However, circadian rhythmicity within the hive follows a more complex pattern, whereby activity peaks are shifted thanks to the spatio-temporal organization of collective division of labor, ensuring that the different tasks in the hive are performed with an appropriate time-shift with respect to the foraging activity. A single mechanism, i.e. mechanical bumping, is sufficient to generate this remarkable organization in the beehive.

## 5 Methods

### Beekeeping

We kept queenright colonies of *Apis mellifera carnica* in a one-frame observation hive at coordinates 52.457130, 13.296285 (Berlin, Germany). Two colonies were recorded - colony A from July 24th to September 19th in 2016 and colony B from July 4th to October 15th in 2019. Data entering our analysis was recorded between August 1st and August 25th (colony A), and between August 20th and September 14th (colony B). Colony A started with ca. 2000 bees, colony B with ca. 1500 bees. New marked bees were added throughout the experiment. The observation hive was located indoors, bees had access to the outside environment through a flexible plastic tube connecting the hive entrance to a hole in the window. A landing board was affixed at the outside to ease the bees’ exit and entrance. Data in Fig. 1, 2 and 3 of the main paper relate to colony B, the corresponding data for colony A are in the Supplementary Information (section 10).

### Adding marked bees

Brood from either the observation colony itself or another colony was kept in an incubator (temperature 34°C). Freshly emerged individuals were removed every day from the brood comb and individually marked at least twice a week. Bees were marked by first removing hair from the thorax using a wet toothpick, applying a thin layer of shellac, and attaching a curved, circular marker showing a binary code (for details see [32]). The number of bees marked per batch varied but never exceeded 156. A record of IDs and their respective marking dates was added to a spreadsheet. We introduced marked bees to the colony through a backdoor entrance. We prevented the emergence of unmarked bees inside the colony by exchanging the comb every 21 days. For colony A, a total of 3166 bees were marked, and throughout the data analysis period we detected 1917 unique individuals. For colony B, a total of 5099 marked bees were introduced, and we detected 3404 unique individuals during the data analysis period. Some marked bees were not accepted by the colony or might have lost their markers. Rejection and marker loss rates may vary from cohort to cohort and were not assessed quantitatively.

### Bee monitoring

The two sides of the observation hive were video recorded with high-resolution cameras (2 x 12 MP per side in colony A, see [32] for details, and one 12 MP camera per side in colony B described in [34]), illuminated by synchronized infrared LED flashes (5 ms light duration) triggered by an Arduino microcontroller. Each comb side was imaged at a rate of 3 Hz for colony A and 6 Hz for colony B, alternating between both sides to avoid backlighting and ensure optimal contrast. Colony B had an additional third camera installed at the far end of the entrance tube providing a 10 Hz recording of bees as they entered or left the hive. All video data was stored to disk for analysis. We then used the BeesBook system [32, 33, 26, 35, 4, 34, 8] to automatically identify and track all marked bees over the whole duration of the recording. This yielded, for each animal and time point: unique bee ID, age, timestamp, planar coordinates, 3-D orientation. The BeesBook computer vision pipeline assigns every detection a confidence score that reflects the system’s uncertainty of decoding the ID and orientation. The trajectory data was transferred to a PostgreSQL database for fast queries in our analyses. To reject false-positive detections for individuals that did not exist, we implemented a Bayesian change point model to calculate the most likely time of death for all individuals in the dataset [34]. Our marking protocol was used to filter out detections with IDs that were not yet in use.

### Extracting trajectories

Sequential detections were linked to tracklets using ID information, orientation and proximity using a learned model, tracklets were linked to trajectories using a second machine learning model, as described elsewhere [4]. The trajectory-level ID and confidence were obtained by median filtering the ID and confidence values for all detections within each trajectory. Bee trajectories with confidence scores lower than 0.1 were excluded. We applied a correction procedure to correct for misalignment of certain bee tag orientation decodings relative to their body orientation as described elsewhere [8]. The resulting trajectories were used to calculate each bee’s movement speed by dividing the Euclidean distance between two consecutive detections by the elapsed time (which was either 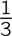 s for colony A or 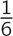 s for colony B, or multiples thereof in case of time points with missing detections). Movement speeds greater than 15 mm/s were considered unrealistic outliers and were removed (leading to 0.1% and 0.2% removed data points in colony A and B respectively).

### Circadian quantification

For each bee, we calculated circadian strength *R*^2^ and phase *ϕ* using the cosinor model [11, 18, 2]. Speeds were subsampled by taking the median for each hour to reduce residual dependence. For each day the fit was calculated using a three-day window (i.e., target day including the preceding and the following day, yielding windows of 72 time points, i.e., hours). We defined the movement speed *v* at a time step *t_i_* (with *i* = 0 *…* 71, and *t_i_* = *−*36 + *i*) as a function of the baseline mean speed (M), the amplitude of the oscillation (A) and the phase (*ϕ*) with a fixed period of *P* = 24h:

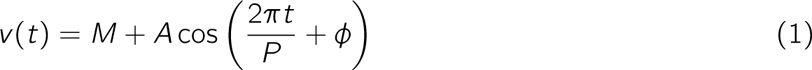

With 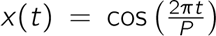 and 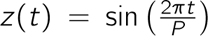 as known variables, and *β* = *A* cos(*ϕ*) and *γ* = *−A* sin(*ϕ*) as unknown parameters, this yields:

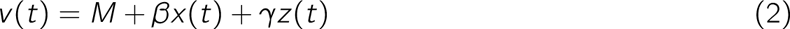

We solved this linear regression problem for *β* and *γ* by using ordinary least squares. Then we deduced the parameters *A* and *ϕ* by calculating 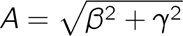 and 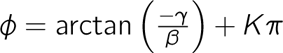 where *K* is an integer [6]. We used an F-test [6] to test for a statistically significant amplitude greater than zero (”zero amplitude test”) to identify rhythmic and non-rhythmic bees. For simplicity, in the figures we convert *ϕ* into the corresponding time of day *T* (*ϕ*) with values between 0 and 24h.

We used the prominence of the fit (*R*^2^) to indicate the strength of the rhythm. We confirmed that *R*^2^ serves as a good estimator for rhythm prominence by correlating it with two other measures of rhythmicity: first, we regressed it against the amplitude *A* (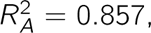 *p <* 0.001, *r* = 0.926 for colony A and 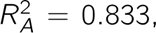 *p <* 0.001, *r* = 0.913 for colony B), and second, against the day-night differences in bee movement speed, as difference of average movement during daytime (9:00-18:00) and night time (21:00-6:00) (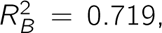 *p <* 0.001, *r* = 0.848 for colony A and 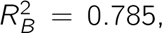 *p <* 0.001, *r* = 0.886 for colony B).

Synchronicity *S*, defined as the phase consistency within groups, was quantified by the standard deviation of the phase (*ϕ*) values across the respective individuals, normalized to [0,1] with *S* = 1 corresponding to the 5%-quantile and *S* = 0 corresponding to the 95%-quantile.

### Physical interactions

We defined two bees to interact when (1) they were detected simultaneously in the hive with a confidence above 0.25, (2) the distance between their thorax markings was less or equal 14 mm, and (3) their body poses were classified as “touching”. To define touching, we modeled both interaction partners as rectangles of 14 *×* 6 mm (approx. the size of a bee), centered on the bee and aligned with her body axis. The masks were then checked for overlap using a logical AND operation. Each interaction was counted as a single event. Since interactions can span multiple frames, subsequent interaction detections between a given pair were considered belonging to the same interaction event if the gaps between them were shorter than one second. For each detected interaction we recorded the position within the nest, timestamp, relative angle of the two bees and overlap area (at the start of the interaction) as well as the duration of the interaction. We treated each individual as a focal bee, yielding two data points for each interaction. For each focal bee we recorded: relative touching positions in the focal bee’s frame of reference, age of the bee, rhythmicity information (*ϕ* and *R*^2^), speed, and change in speed after the interaction. The speed change was calculated as the average speed in a 30-seconds window after the interaction minus the average speed during 30 seconds before the interaction started.

### Statistical analysis of physical interactions

To assess whether interactions yield speed changes that are significantly different from non-interacting bees, we defined a null model. For each real interaction, we selected two random bees present in the nest at the same time points and calculated the speed changes resulting from these (fictive) interactions. We analyzed median speed change, movement speed at the start of the interaction, phase, and prominence of rhythm. For each of these we divided the focal and non-focal bees into six quantiles resulting in 36 combinations. Measured interactions and the null model were compared using Welch’s t-test.

### Body-centered physical interactions

To quantify speed change as a function of the relative touching positions we defined an area of 16 *×* 8 mm representing the focal bee (plus 1 mm padding) and discretized this area into 16 *×* 8 bins (yielding a spatial resolution of 1 mm / bin). For every given interaction, the speed change values of both the focal bee and her interaction partner were accumulated in the focal bee’s bins corresponding to its intersection with the non-focal bee’s rectangle. Fine-grained analysis included segmenting the data for different combinations of criteria, such as duration of the interaction or characteristics of the participating bees, e.g. age.

### Agent-based modeling

We created an agent-based simulation to study the effects of speed transfers from circadian agents to non-circadian ones. The spatial arrangement mirrored the observation hive: a rectangular, two-dimensional plane bounded by walls on each side. We defined two agent populations: one representing older foraging bees, clustered at one side of the nest, and with a prominent circadian rhythm, the second representing younger nurse bees, concentrated in the middle of the nest, and without a circadian rhythm. Each agent was modeled as a point with a given 2-D position and orientation. For every simulation step, agents moved into the direction they were facing. The amount of movement (i.e., the speed) was determined by three components. Every bee received a baseline speed drawn from a Gaussian random distribution (clipped to 0 to avoid negative speeds). The forager group received an additional sinusoidal driver representing periodic speed changes. For interacting bees the slower bee received the speed difference with the faster bee. Interactions were determined by thresholding inter-individual distance. After taking a step, the orientation of each agent was changed by adding a Gaussian random sample and additionally both groups were pulled back towards the center of their respective initial distributions. Agents that reached the boundary of the arena were reflected back. The resulting motion data was analyzed with the same procedure as the biological observations.

### Speed transfer cascade trees

Binary interactions create a tree-like interaction history that can be traced, such as bee a being activated by bee b1 and bee b2, which in turn are activated by bees c1..cn, and so on. Interactions were only considered between 10:00 and 15:00. We selected a random subset of *n* = 1000 bees from those that were significantly rhythmic, younger than 5 days old, and reached their peak activity after 12:00. These bees were taken as our target group for the ends of interaction chains. We then identified, for each bee, all interactions that caused a positive change in speed, and traced back interactions in time. This yielded a tree structure, where the root vertex was fixed to be a young rhythmic bee and all bees that had activated her were added in the previous timestep. The maximum time window between two interactions was set to 30 minutes, and the maximum duration of a cascade was set to be 2 hours. After creating such trees, we extracted all paths from the root back to the leaf nodes. To test if phase *ϕ*, *R*^2^, or age differed between the leaf nodes and the general hive we used a Brunner-Munzel test [5].

## 6 Data Availability

The raw data (video files, trajectories) are approximately 400 TB in size and available from the corresponding author on reasonable request. All derived datasets (speeds, interactions, etc) analysed during the current study are available online: https://github.com/BioroboticsLab/speedtransfer.

## 7 Code Availability

All code is available online: https://github.com/BioroboticsLab/speedtransfer.

## 8 Author Contributions

Conceptualization: J.M., W.K., B.W., D.M.D., A.Z., M.L.S., C.G.G. and T.L.; Methodology: J.M., W.K., B.W., D.M.D., A.Z. and T.L.; Software: J.M., W.K., B.W., D.M.D., A.Z. and T.L.; Resources, supervision: T.L.; Project administration: T.L.; Data curation: J.M., W.K., B.W. and D.M.D.; Writing: J.M., W.K., B.W., D.M.D., M.L.S., C.G.G and T.L; Visualization: J.M., W.K., B.W. and D.M.D.

## 10 Supplementary Information

### Figure S1: Relates to Fig. 1. Circadian rhythms across bee age and nest space for colony A. While Fig. 1 in the main manuscript relates to colony B, here we show the same analysis for colony A

**Figure S1a:**
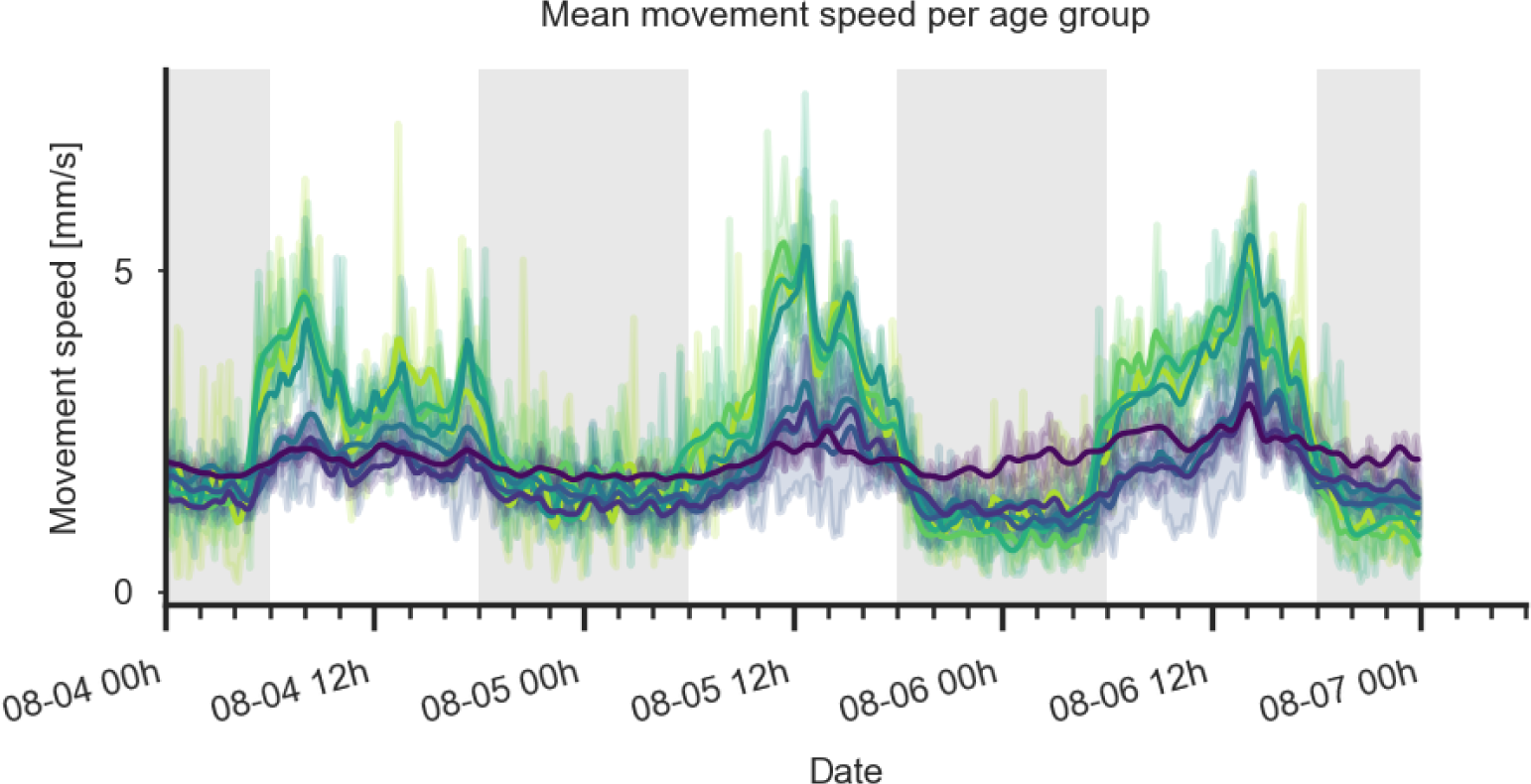
All age groups show higher speeds during the day than at night, but the difference increases with bee age. Example movement speed [mm/s] of bees for three days, split by age from young (purple) to old (yellow). Data was binned to hours, and smoothed with a Gaussian 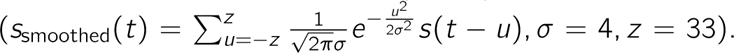 Lighter bands show 95% confidence intervals of the age group unsmoothed velocities.

**Figure S1c:**
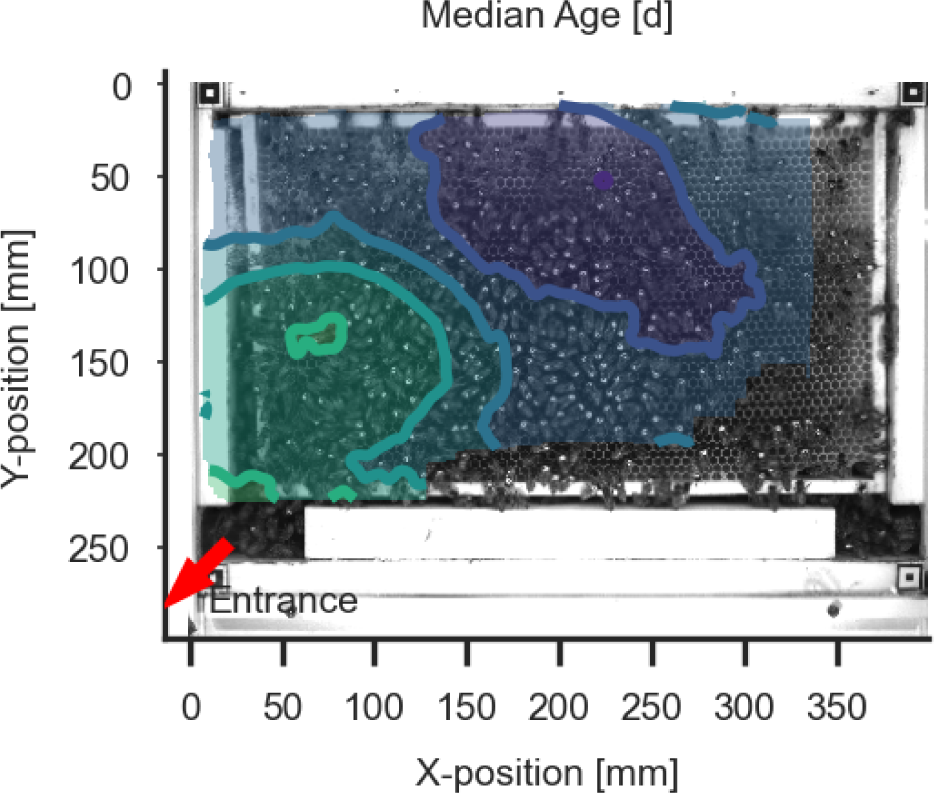
Spatial distribution of median age over the comb. Older bees (foragers) tend to be closer to the entrance, while younger bees (nurses) are more concentrated in the center of the comb.

**Figure S1d:**
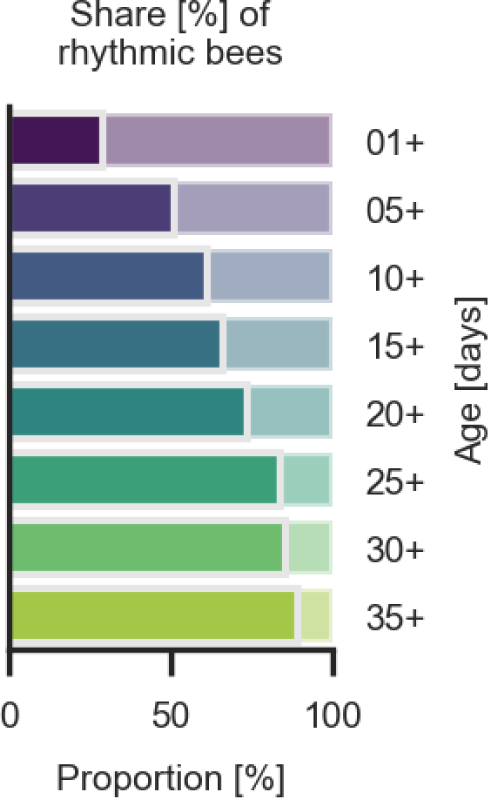
With increasing age, the share of significantly circadian bees increases. Bees are split in age groups as in (a), and the proportion of significantly rhythmic bees is shown for each group.

**Figure S1e:**
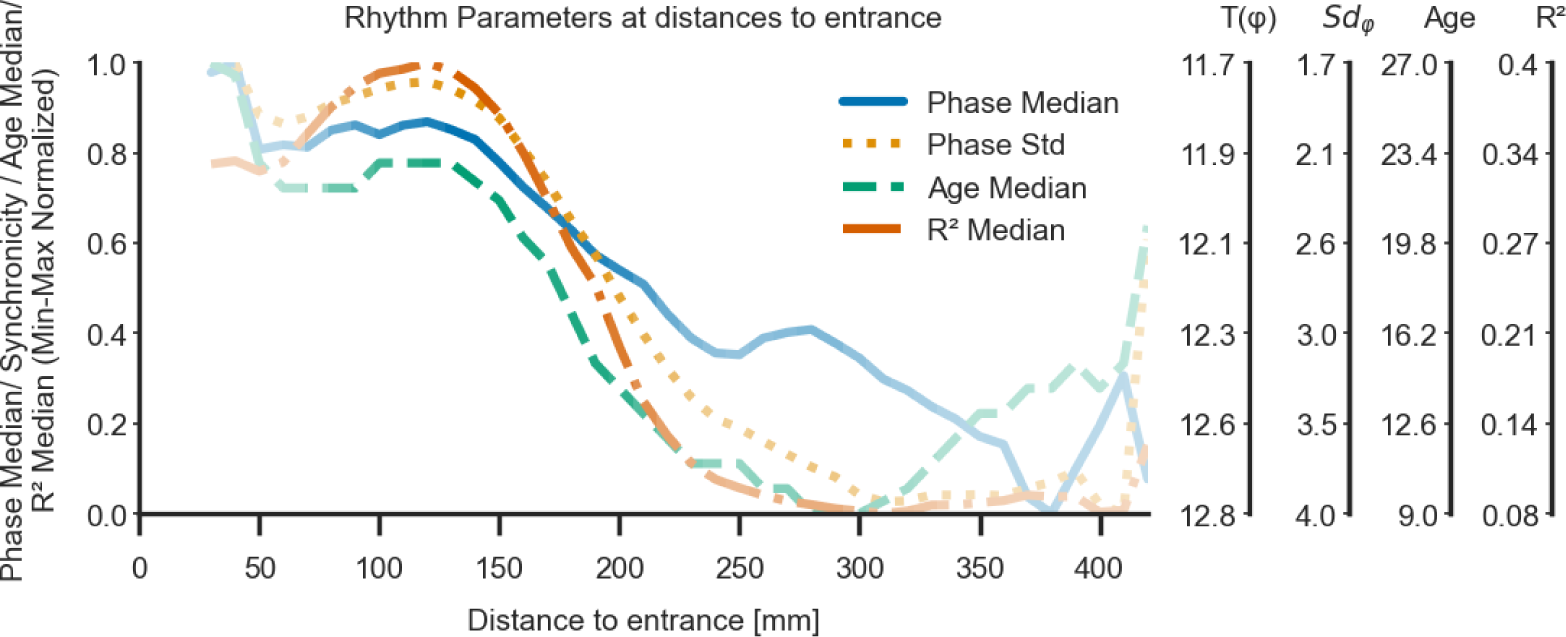
Circadian parameters decrease with increasing distance from the entrance. Curves indicate peak phase *P* (daytime of maximum), phase standard deviation (hours), and circadian prominence *R*^2^ as calculated with the cosinor fit, relative to distance to entrance. Bee age (days) also decreases with increasing distance from the entrance. Curve color saturation indicates data density (i.e., how many bees were in that area). Synchronicity is defined as the standard deviation of the phase. The four scales to the right correspond to the original values for the four normalized traces.

### 10.1 Figure S2. Relates to Fig. 2. Dynamic physical interaction leads to speed transfer across bees (colony A). While Fig. 2 in the main manuscript relates to colony B, here we show the same analysis for colony A

**Figure S2:**
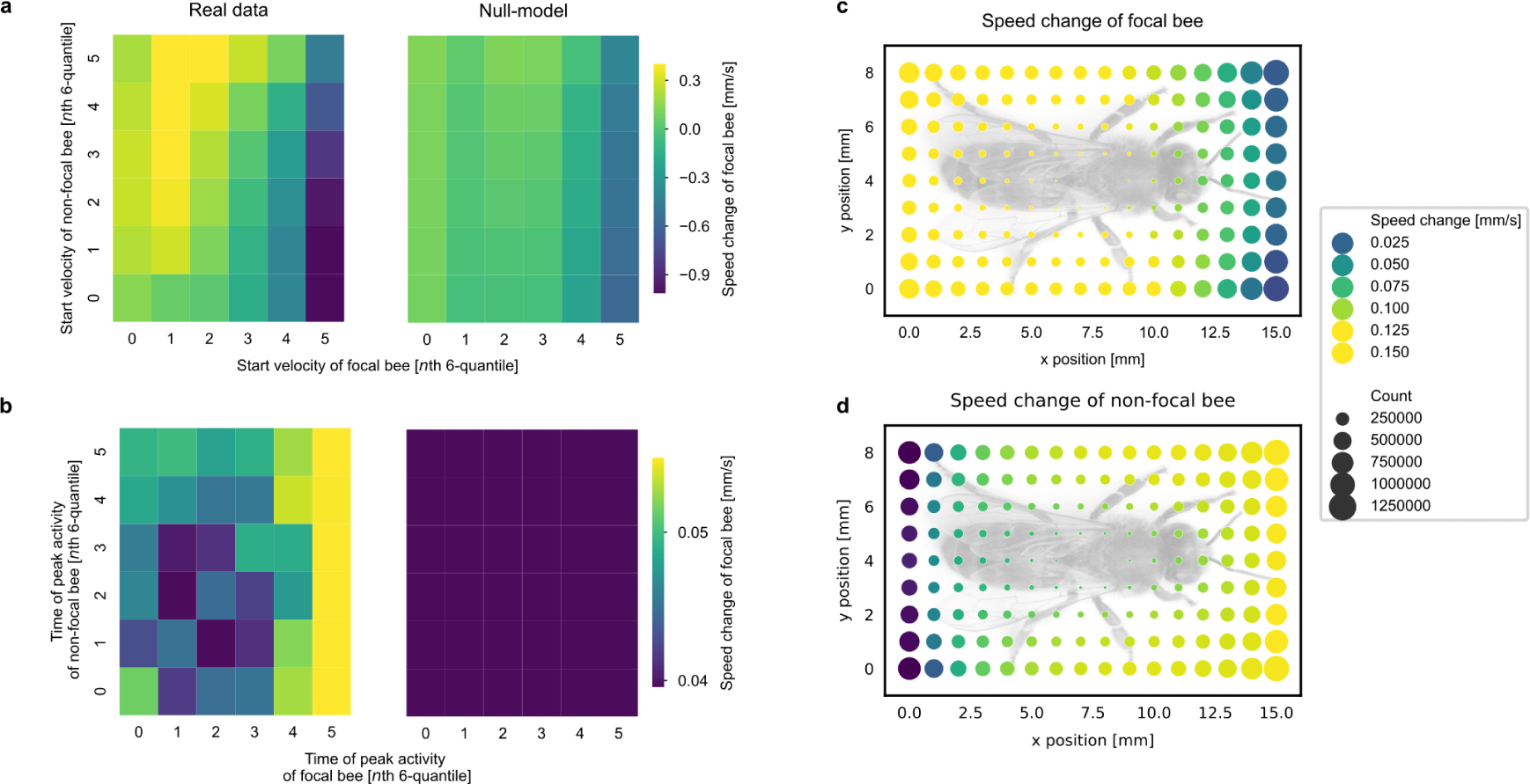
**(a)** Fast bees are slowed down when bumping into slow bees. Focal bees (x-axis) and non-focal bees (y-axis) were sorted into 6 speed quantiles. For each combination, the speed change of the focal bee was color coded: speed decreases (blue hues) were particularly prominent for previously fast bees (6th quantile). The right panel shows data from a randomly shuffled null-model (see methods), experimental data is significantly different from the shuffled distribution (see Supplementary Note 2 for statistical values). **(b)** Circadian phase *ϕ* of the interaction partner influences speed transfer. Focal and non-focal bees were binned in 6 quantiles depending on the phase of their cosinor fitted circadian activities. Focal bees with late phase led to higher increases in non-focal bees. The randomly shuffled null-model (right panel) showed no phase effect. **(c)** Bees are slowed down when bumping into another bee from the front, and accelerated when bumped from the back. Image shows speed change of the focal bee (shown schematically in grey) relative to the interaction location along her body. Speed changes resulting from interacting at the front are negative (blue hue), interactions at the back are positive (yellow hue). **(d)** Bees are slowed down when bumping into the back of another bee, and accelerated when hit by a head. Image shows speed change of the non-focal bee relative to the interaction location along the body of the focal bee (shown schematically in grey). Speed changes are negative along the back of the focal bee, and positive along her front. In (c) and (d), circle sizes indicate the number of observations in each point.

### 10.2 Local interactions yield speed transfer: P-values for testing interaction metrics against, non-interaction null-model

For any of the two colonies, focal bees (columns) and non-focal bees (rows) of all interactions were grouped into six quantiles of four features (speed at interaction start, phase, *R*^2^, age). For each combination, the change in velocity of the focal bee was compared to a null model (see Methods) using a Welch test. The tables show the corresponding p-values. Tests were performed on independent subsets of data. Tables 1-4 relate to colony B, tables 5-8 relate to colony A.

**Supplementary Table 1:** P-values of Welch tests comparing the speed changes of bees grouped into 6 *×* 6 pairings reflecting their respective interaction start speed quantiles (colony B). Relates to Fig. 2a.

|  | 1 | 2 | 3 | 4 | 5 | 6 |
| --- | --- | --- | --- | --- | --- | --- |
| 1 | 0.0 | 0.0 | 0.0 | 0.0 | 0.0 | 8.52e-06 |
| 2 | 4.33e-22 | 1.11e-176 | 0.0 | 3.39e-244 | 0.02 | 0.0 |
| 3 | 6.09e-17 | 3.57e-74 | 1.90e-181 | 1.56e-44 | 7.29e-38 | 0.0 |
| 4 | 0.81 | 1.03e-16 | 2.18e-08 | 1.11e-08 | 3.16e-154 | 0.0 |
| 5 | 0.0 | 0.0 | 0.0 | 0.0 | 0.0 | 0.0 |
| 6 | 0.0 | 0.0 | 0.0 | 0.0 | 0.0 | 0.0 |

**Supplementary Table 2:** P-values of Welch tests comparing the speed changes of bees grouped into 6 *×* 6 pairings reflecting their respective phase quantiles (colony B). Relates to Fig. 2b.

|  | 1 | 2 | 3 | 4 | 5 | 6 |
| --- | --- | --- | --- | --- | --- | --- |
| 1 | 4.42e-91 | 3.94e-64 | 3.84e-71 | 5.82e-76 | 9.68e-52 | 9.39e-168 |
| 2 | 2.35e-125 | 2.53e-158 | 9.99e-180 | 4.27e-248 | 1.77e-298 | 3.44e-134 |
| 3 | 2.51e-135 | 5.50e-149 | 3.05e-180 | 1.86e-290 | 7.84e-214 | 7.56e-136 |
| 4 | 1.39e-122 | 7.62e-118 | 7.88e-151 | 3.44e-147 | 7.15e-126 | 1.23e-137 |
| 5 | 2.53e-122 | 8.93e-139 | 1.26e-81 | 3.61e-103 | 2.47e-71 | 2.38e-134 |
| 6 | 3.48e-150 | 2.66e-107 | 2.41e-92 | 1.28e-73 | 8.88e-49 | 3.82e-120 |

**Supplementary Table 3:**
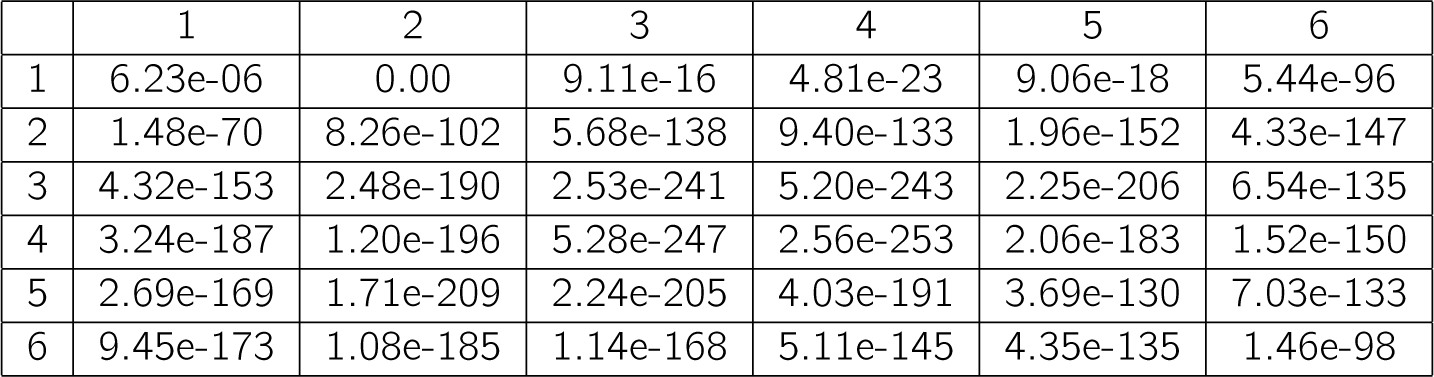
P-values of Welch tests comparing the speed changes of bees grouped into 6 *×* 6 pairings reflecting their respective *R*^2^ quantiles (colony B).

**Supplementary Table 4:** P-values of Welch tests comparing the speed changes of bees grouped into 6 *×* 6 pairings reflecting their respective age quantiles (colony B).

|  | 1 | 2 | 3 | 4 | 5 | 6 |
| --- | --- | --- | --- | --- | --- | --- |
| 1 | 4.11e-67 | 7.07e-47 | 4.92e-25 | 1.89e-13 | 1.37e-18 | 5.69e-47 |
| 2 | 8.55e-152 | 8.63e-121 | 1.57e-84 | 1.06e-69 | 2.62e-77 | 3.85e-58 |
| 3 | 2.70e-180 | 6.01e-184 | 4.82e-184 | 6.26e-131 | 6.95e-65 | 4.58e-56 |
| 4 | 2.36e-258 | 1.06e-271 | 2.19e-255 | 7.13e-121 | 1.82e-81 | 3.86e-75 |
| 5 | 0.0 | 0.0 | 4.09e-151 | 7.35e-84 | 6.79e-61 | 2.97e-58 |
| 6 | 0.0 | 0.0 | 1.86e-80 | 5.60e-39 | 4.08e-35 | 1.76e-17 |

**Supplementary Table 5:** P-values of Welch tests comparing the speed changes of bees grouped into 6 *×* 6 pairings reflecting their respective interaction start speed quantiles (colony A).

|  | 1 | 2 | 3 | 4 | 5 | 6 |
| --- | --- | --- | --- | --- | --- | --- |
| 1 | 9.61e-192 | 0.0 | 1.12e-300 | 3.85e-309 | 5.21e-284 | 0.0 |
| 2 | 0.0 | 5.07e-137 | 1.82e-18 | 0.10 | 0.15 | 0.05 |
| 3 | 2.64e-157 | 1.15e-134 | 1.44e-79 | 1.62e-124 | 5.86e-122 | 1.40e-70 |
| 4 | 3.78e-299 | 0.0 | 0.0 | 0.0 | 0.0 | 0.0 |
| 5 | 0.0 | 0.0 | 0.0 | 0.0 | 0.0 | 0.0 |
| 6 | 0.0 | 0.0 | 0.0 | 0.0 | 0.0 | 0.0 |

**Supplementary Table 6:**
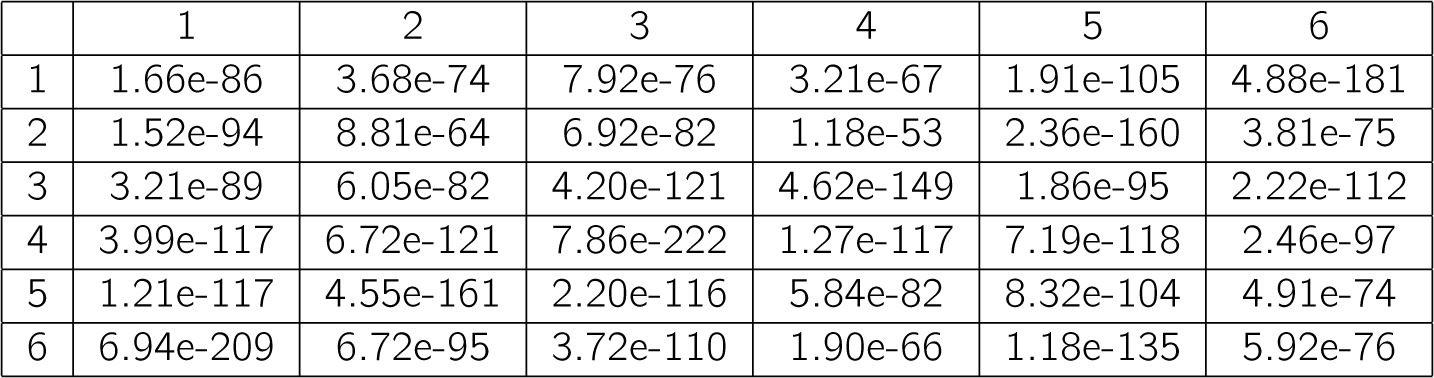
P-values of Welch tests comparing the speed changes of bees grouped into 6 *×* 6 pairings reflecting their respective phase quantiles (colony A).

**Supplementary Table 7:** P-values of Welch tests comparing the speed changes of bees grouped into 6 *×* 6 pairings reflecting their respective *R*^2^ quantiles (colony A).

|  | 1 | 2 | 3 | 4 | 5 | 6 |
| --- | --- | --- | --- | --- | --- | --- |
| 1 | 1.32e-23 | 6.48e-34 | 1.58e-37 | 3.70e-54 | 2.58e-123 | 5.85e-282 |
| 2 | 3.00e-81 | 1.19e-90 | 1.55e-94 | 4.80e-109 | 2.56e-205 | 2.87e-148 |
| 3 | 2.23e-102 | 1.17e-101 | 6.44e-116 | 3.43e-188 | 1.77e-167 | 2.05e-92 |
| 4 | 3.96e-93 | 5.44e-100 | 2.36e-171 | 1.51e-130 | 5.61e-128 | 1.08e-69 |
| 5 | 5.14e-102 | 2.68e-157 | 8.98e-99 | 1.52e-114 | 6.64e-95 | 1.56e-77 |
| 6 | 3.05e-152 | 1.70e-94 | 4.49e-91 | 1.93e-85 | 1.15e-73 | 1.82e-59 |

**Supplementary Table 8:** P-values of Welch tests comparing the speed changes of bees grouped into 6 *×* 6 pairings reflecting their respective age quantiles (colony A).

|  | 1 | 2 | 3 | 4 | 5 | 6 |
| --- | --- | --- | --- | --- | --- | --- |
| 1 | 3.69e-37 | 3.43e-70 | 1.42e-85 | 6.38e-78 | 4.35e-120 | 3.12e-266 |
| 2 | 1.91e-76 | 1.16e-69 | 5.82e-72 | 1.15e-55 | 1.21e-129 | 6.208e-127 |
| 3 | 2.19e-102 | 9.80e-87 | 1.41e-51 | 1.78e-147 | 1.38e-64 | 6.25e-112 |
| 4 | 3.41e-104 | 1.94e-73 | 1.64e-175 | 3.70e-41 | 3.38e-70 | 1.30e-73 |
| 5 | 6.30e-206 | 0.0 | 3.47e-56 | 2.58e-66 | 1.33e-52 | 4.00e-58 |
| 6 | 0.0 | 5.10e-152 | 1.46e-45 | 1.29e-35 | 3.57e-19 | 5.09e-46 |

### 10.3 Speed transferred depends on attributes of interacting bees

**Figure S3:**
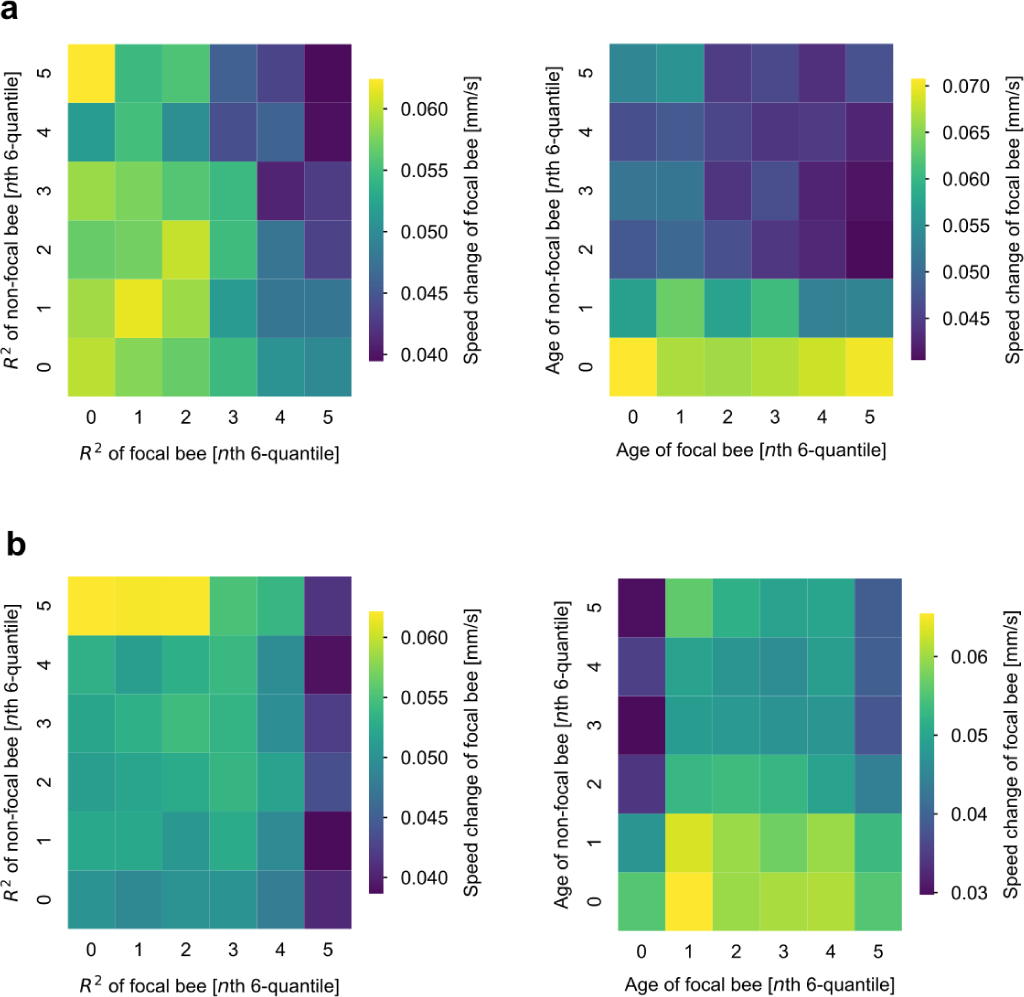
Average speed change of focal bee for respective combinations of 6 *×* 6 quantiles of *R*^2^ and age for colony A (row a) and colony B (row b), respectively.

### 10.4 Resting bees: speed transferred irrespective of where focal bee was touched

**Figure S3:**
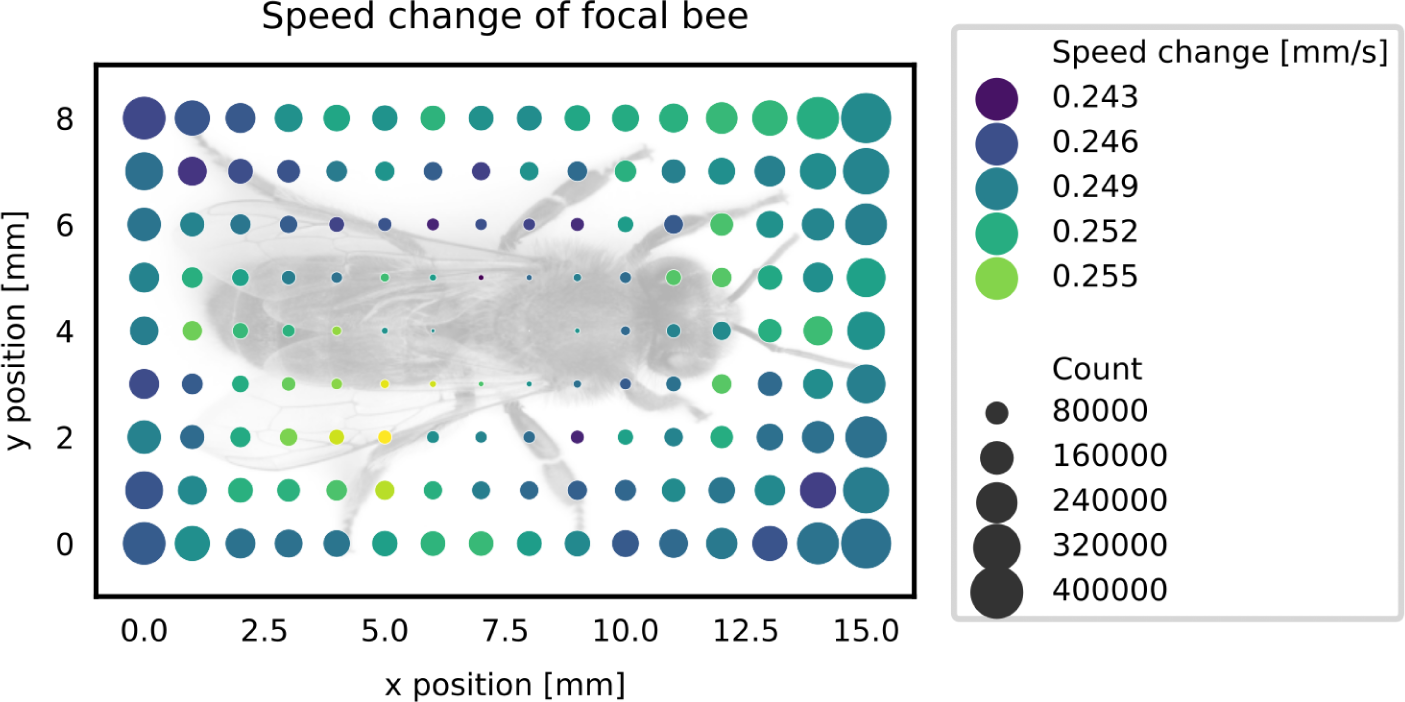
Resting bees are activated irrespective of where they touch their interaction partner. The average speed changes after an interaction for bins of 1*×*1 mm show positive values consistently around the focal bee’s body.

